# Dissociable neuronal substrates for positive and negative valence stimuli in the nucleus accumbens

**DOI:** 10.64898/2026.01.24.701496

**Authors:** Raquel Correia, Ana Verónica Domingues, Leandro AA Aguiar, Margarida Cunha-Macedo, Daniela Vilasboas-Campos, Natacha Vieitas-Gaspar, Rafael M. Jungmann, Eduardo Teixeira, Bárbara Coimbra, Ana João Rodrigues, Carina Soares-Cunha

## Abstract

The nucleus accumbens (NAc) responds to both natural and artificial rewards and to aversive stimuli; however, it remains unclear whether these opposing valence signals engage distinct neuronal ensembles. Here we used Fos-CreERT^2^-based activity-dependent tagging to label NAc neuronal ensembles activated by cocaine or foot shock. We found that cocaine ensemble consisted predominantly of dopamine D1 receptor-expressing medium spiny neurons (D1-MSNs), whereas foot shock ensemble similarly recruited D1- and D2-MSNs. One-photon calcium imaging in freely moving mice revealed that acute cocaine primarily excited D1-MSNs while inhibiting the majority of D2-MSNs, whereas foot shock induced excitatory responses in both types of MSNs. Optogenetic reactivation of the cocaine-ensemble elicited a strong behavioural preference, whereas reactivation of the shock-ensemble produced no significant behavioural effect.

Together, these findings demonstrate that cocaine recruits a functionally specific NAc ensemble distinct from that recruited by shock, providing mechanistic insight into the valence-specific neuronal substrates underlying reward and aversion processing.

## Introduction

Reward- or aversive-based learning represents a conserved evolutionary survival strategy across species^1^. In nature, some stimuli are inherently appetitive or aversive, leading to approach or avoidance, respectively^2^. In addition, individuals must learn to associate external cues with emotionally relevant outcomes to guide behavior. Importantly, disruptions in these stimuli-responses or cue-outcome associations emerge in several psychiatric disorders, namely drug addiction and depression^3,4^.

The NAc plays a central role in encoding positive and negative valence stimuli^5^. This striatal region receives dense dopaminergic innervation from the ventral tegmental area (VTA), and transmits information related to salience, value, context, and motor planning onto basal ganglia outputs^6^. These signals are sent onto downstream targets by GABAergic MSNs that express either dopamine receptor D1 or D2, or both^6^. Earlier optogenetic studies have proposed that these two neuronal populations had opposing behavioral roles, with D1-MSNs mediating reward and D2-MSNs aversion^7,8^. However, recent evidence shows a complementary role in valence-related behaviors^9–16^. We showed that optical activation of D1- or D2-MSNs enhances cue-driven motivation for food^9,15^ and optogenetic activation of NAc core D1- or D2-MSNs drives reward or aversion, depending on specific activity patterns^14^. Interestingly, brief activation of either population increased cocaine conditioning, whereas prolonged imaging revealed that both D1- and D2-populations encode positive or negative stimuli^13,16,17^. Curiously, no clear cell specificity in response to sucrose or shock was reported, since around 70% of all recorded neurons responded to both stimuli^13^. This overlap is in opposition to basolateral amygdala (BLA), where a clear segregation of valence-encoding neurons is observed^18,19^.

Given its importance within the reward circuit, the NAc also responds to drugs of abuse^20–23^. Population D2-MSN (but not D1) activation decreased cocaine conditioning^14^. Furthermore, single cell calcium activity of D1- and D2-MSNs shows similar patterns during a cocaine-conditioned place preference (CPP) test^24^. In addition, D1-MSNs exhibit stable, time-locked activity during cocaine-seeking, but not in sucrose-seeking^25^. These and other studies thus suggest differential engagement of D1- and D2-MSNs in reward processing and support the notion of specific ensembles encoding natural- or drug-related responses^21,25,26^.

Despite advances, it is still unclear if cocaine activates a “generalized appetitive” or a drug-specific ensemble. In addition, the identity of an “aversive ensemble” is not characterized in detail, nor we know the degree of overlap with the cocaine ensemble. To address these gaps, we employed a Fos-based labeling approach to permanently tag the neuronal ensembles activated by cocaine (appetitive non-natural stimulus) and foot shock (aversive stimulus)^27–30^. We demonstrate that cocaine and foot shock recruit partially overlapping but functionally distinct accumbal ensembles, with cocaine-ensemble uniquely capable of driving behavioral preference. These findings provide novel insights into the neural encoding of drugs and aversive events, highlighting a functional dissociation in the behavioral impact of reactivating stimulus-specific neuronal ensembles in the NAc.

## Results

### Acute cocaine exposure recruits a D1-MSN-enriched ensemble in the anterior NAc

To determine the anatomical distribution of the NAc ensemble that represents acute cocaine exposure, we used the TRAP2;Ai14 transgenic mouse line^30^, that relies on the c-fos immediate early gene locus to drive the expression of tamoxifen-inducible CreER.

Mice were habituated for 7 days to intraperitoneal (i.p.) injections of saline (vehicle). On the test day (day 8), mice received i.p. injections of either saline or cocaine hydrochloride (20mg/kg) immediately followed by 4-hydroxitamoxifen (4-OHT) i.p. administration (Figure 1A). Subsequent RNAscope® *in situ* hybridization with probes for Drd1, A2A and tdTomato (*A2a* was used as a marker for D2-MSNs^31–33^) (Figure 1C) revealed that the cocaine ensemble was significantly larger than the saline ensemble (Figure 1D). Specifically, a larger proportion of tdTomato neurons was found in the NAc core (NAcC) and lateral shell (latNAcSh), and a trend for increased numbers in the dorsomedial shell (dmNAcSh) and ventromedial shell (vmNAcSh). The cocaine ensemble presented a specific anteroposterior (AP) distribution (Figure 1E; saline, Figure S1B), with highest proportion of cocaine-activated cells at the more anterior portions.

**Figure 1.**
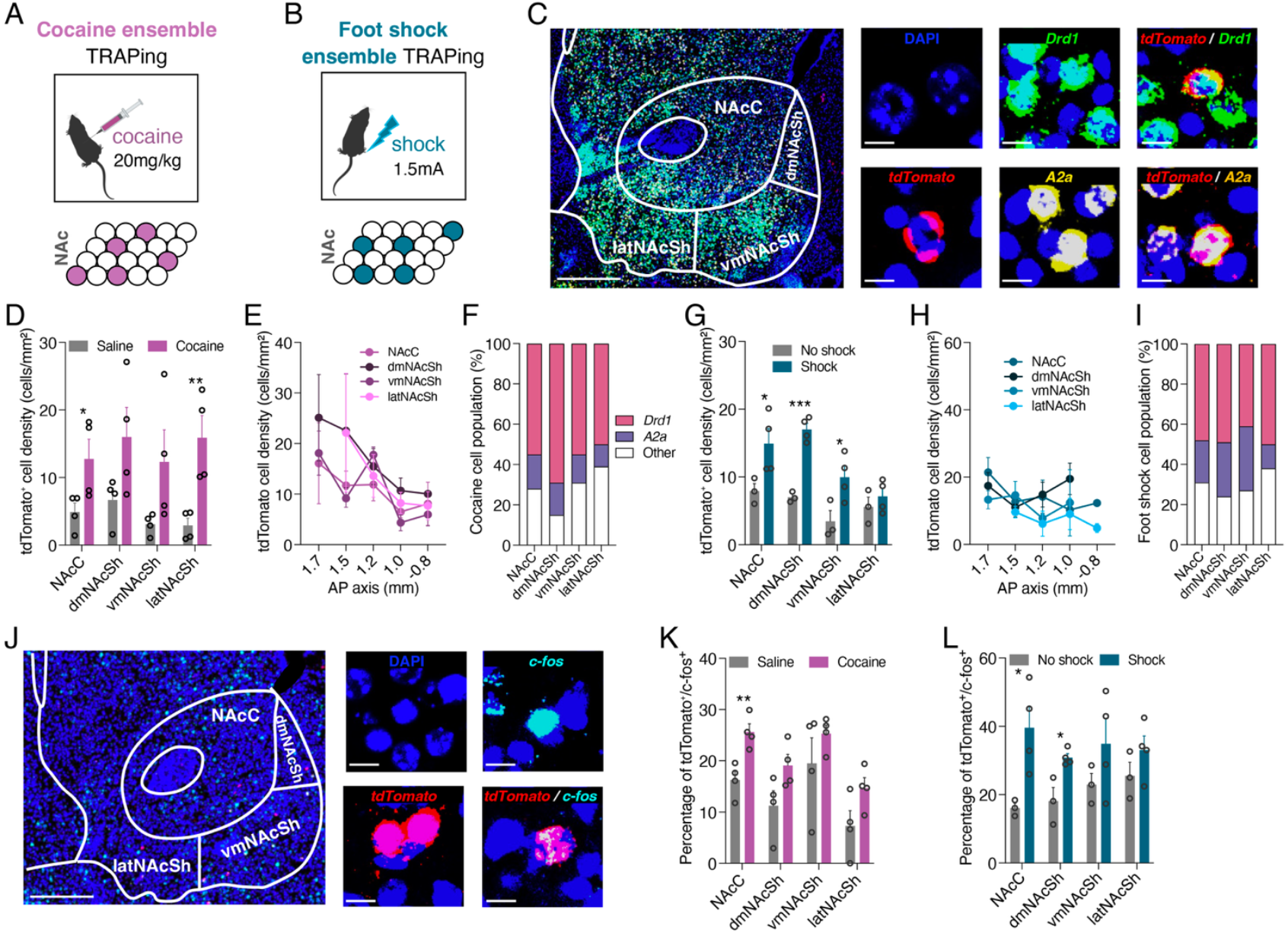
Distinct NAc ensembles are recruited by acute cocaine and foot shock exposure. **A** After 7 days of habituation to saline i.p. injections, TRAP2;Ai14 mice received a single i.p. injection of cocaine (control mice received saline) and i.p. injection of 4-OHT. **B** After 7 days of habituation to the behavioural box and saline i.p. injections, TRAP2;Ai14 mice received 1 session of foot shock delivery (20-foot shocks, 1.5 mA, 1s; 30s interval) followed by 4-OHT i.p. administration. **C** Representative images for *tdTomato* (tdTom+), *Drd1* (D1-MSN+) and *A2a* (D2-MSN+) mRNA; DAPI (blue) marks nuclei; scale bar=500µm; inset scale bar=10µm. **D** Cocaine administration recruited a significantly larger ensemble than saline exposure in NAcC (left; n_cocaine_=4, n_saline_=4; *unpaired t-test, p*=0.0500) and latNAcSh (*p*=0.0095). **E** AP distribution of tdTomato+ neurons across NAc subregions for cocaine-TRAPed ensembles. **F** tdTomato+ cell-type composition within cocaine ensembles. **G** Foot shock delivery recruited a significantly larger ensemble than no shock group in NAcC (right; n_shock_=4, n_no shock_=3), *p*=0.0425), dmNAcSh (*p*=0.0003) and vmNAcSh (*p*=0.0402). **H** Distribution of tdTomato+ neurons across NAc subregions for foot shock-TRAPed ensembles. **I** tdTomato+ cell-type composition within foot shock ensembles. **J** Representative images for *tdTomato* and *c-fos* mRNA; DAPI (blue) marks nuclei; scale bar=500µm; inset scale bar=10µm. **K** Percentage of *tdTomato*/*c-fos* cells of cocaine -> shock mice (NAcC: *p*=0.0067). **L** Percentage of *tdTomato*/*c-fos* cells of shock -> cocaine mice (NAcC: *p*=0.0266; dmNAcSh: *p*=0.0165). Data represent mean ± SEM. \**p*<0.05, *\*\*p*<0.01.

We found that 57% of cells recruited in the cocaine-seeking ensemble were D1-MSNs (Figure 1F) and 15% were D2-MSNs. The saline ensemble presented similar percentage of recruited D1-MSNs (56%), but a higher proportion of D2-activated neurons (31%) (Figure S1A).

All together, these results show that acute cocaine exposure activated a predominantly D1-MSN-enriched ensemble within the NAc and showing an anterior-biased distribution.

### Foot shock recruits D1- and D2-MSNs similarly distributed across medial NAc

Given that accumbal D1- and D2-MSNs significantly respond to unexpected shock delivery^13^, we next mapped and identified the cells of the foot shock-ensemble. TRAP2;Ai14 mice were habituated to the behavior chamber for 10 minutes and to i.p. injections, for 7 days. On the 8^th^ day, mice were placed in the behavior chamber and received 20-foot shocks (1.5mA, 1 second), with an interval of 30 seconds, followed by administration of 4-OHT (Figure 1A). Foot shock recruited more cells than the no shock group (Figure 1G). The shock ensemble exhibited uniform distribution along the anteroposterior axis with no clear anatomical segregation between subregions (Figure 1H). Unlike the cocaine ensemble, the shock ensemble showed proportional recruitment of D1- and D2-MSNs (47 and 32%, respectively; Figure 1I).

These results show that foot shock ensemble is characterized by balanced D1- and D2-MSN recruitment and no clear anatomical segregation.

### Cocaine and foot shock ensembles show significant overlap

To determine whether cells that are activated by a specific stimulus also respond to the stimulus with opposite valence, TRAP;Ai14 mice previously exposed to one of the stimuli (cocaine or shock) were let to recover for 2 weeks and after that period were submitted to the opposite stimulus and endogenous c-fos activation was assessed. RNAscope was performed against tdTomato (first stimulus) and c-fos(second stimulus) (Figure 1J). 15-25% of the cells in the NAc (sub-region dependent) initially activated by cocaine were activated by foot shock (Figure 1K). A significantly higher proportion of activated cells in the NAcC and a trend for increased proportion of activated cells by the second stimulus in the dmNAcSh and latNAcSh in comparison to the respective control group (saline followed by no foot shock) was found (Figure 1K). In mice initially exposed to foot shock, 31-40% of the cells (sub-region depedent) were activated by cocaine administration (Figure 1L). A significantly higher proportion of activated cells by the second stimulus in the NAcC and dmNAcSh in comparison with the control group (foot shock followed by saline) was found (Figure 1L).

### D1- and D2-MSNs activity in response to acute cocaine and unexpected foot shock delivery

We next used microendoscopic calcium imaging in either population within the NAcC and medial shell (mNAcSh) since these areas showed the highest overlap in TRAPed cells between the two conditions. D1- or A2A-cre mice were injected with an adeno-associated virus (AAV) carrying cre-dependent expression of GCaMP6f, followed gradient index (GRIN) lens implantation (Figure 2A-B; Fig. S2A-B). Six weeks later, neural dynamics was recorded in response to a single dose of cocaine (20 mg/kg) and a single session of foot shock in the same animal (Figure 2C-D).

**Figure 2.**
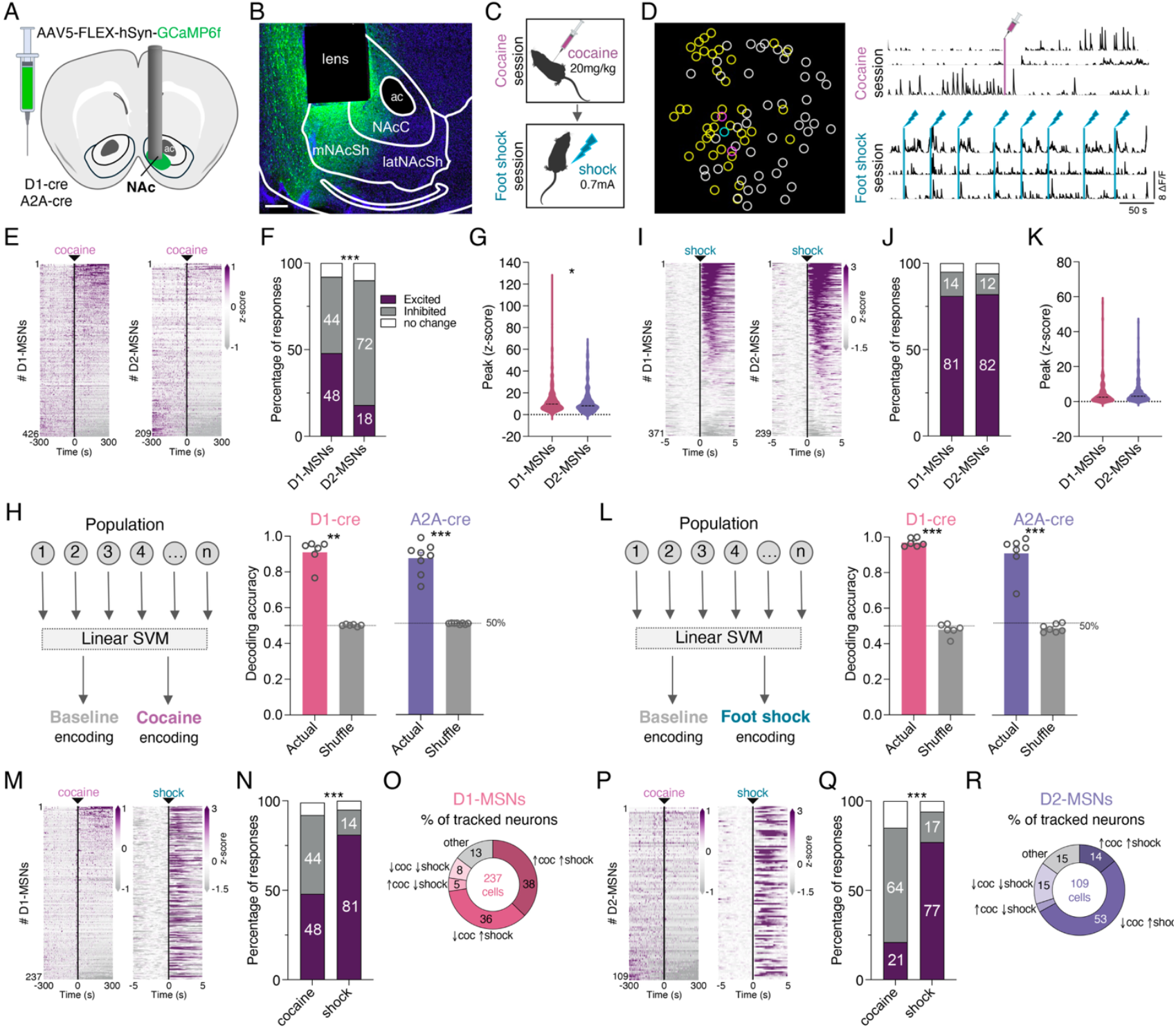
D1- and D2-MSNs display distinct activity in response to cocaine and foot shock. **A** Scheme of calcium imaging recordings in D1- and D2-MSNs. **B** Representative expression of GCaMP6f, depicting GRIN lens location; scale bar=200μm. **C** Schematic representation of cocaine and foot shock exposure. **D** Field of view of an example D1-cre mouse showing neurons responsive to cocaine (pink), to foot shock (blue) and to both (yellow); traces of example tracked single neurons responsive in both sessions; vertical lines in traces represent events. Activity heatmaps of individual D1- and D2-MSNs aligned to **E** cocaine administration (n_D1-MSNs_=426 cells, n_D2-MSNs_=209 cells) or to **I** foot shock delivery (n_D1-MSNs_=372 cells, n_D2-MSNs_=239 cells). Percentage of excited (purple), inhibited (gray) and non-responsive (white) D1- and D2-MSNs in response to **F** cocaine administration or **J** foot shock delivery; single cocaine administration generated significantly more excitation in D1-MSNs than in D2-MSNs (*xi-square test, p*<0.0001). Peak z-score quantification of individual D1- and D2-MSNs after **G** cocaine administration (higher response magnitude in D1-MSNs; *Mann Whitney test*, U=39907, *p*=0.0338), or after **K** foot shock delivery. SVM decoder accuracy for predicting either **H** cocaine exposure (D1-cre, *p*=0.0022; A2A-cre, *p*=0.0002) or **L** foot shock delivery (D1-cre, *unpaired t-test, p*<0.0001; A2A-cre, *Mann-Whitney, p*=0.0006), based on D1- or D2-MSN population activity. Activity heatmaps of individual **M** D1-MSNs (n=237 cells) or **P** D2-MSNs (n=109 cells) tracked over the cocaine administration session and the foot shock delivery session. Percentage of excited, inhibited and non-responsive **N** D1-MSNs or **Q** D2-MSNs in response to cocaine administration or foot shock delivery. Percentage of each type of response of neurons tracked during the cocaine session and foot shock delivery session for **O** D1-MSNs and **R** D2-MSNs. Data represent mean ± SEM. \*\*\**p*<0.001.

Before imaging cocaine-responsive neurons, mice were first habituated for three consecutive days to the open-field arena and to a dummy miniscope. During each habituation session, mice received an i.p. injection of saline at the 5-minute point and remained in the arena for an additional period of 20 minutes. On the day preceding cocaine administration, calcium activity was recorded using the miniscope while mice explored the same arena for 25 minutes; mice received an i.p. saline injection 5 minutes after the onset of the recording. On the following day, mice were again imaged in the same open field and received an i.p. injection of cocaine hydrochloride (20 mg/kg) at the 5-minute point. Cocaine significantly increased the locomotor activity of mice in comparison with the saline session (Figure S2C-D). Individual neurons’ activity was classified as responsive when their activity changed significantly in the 5 minutes after cocaine/saline injection versus baseline (*p*<0.05, Permutation test).

Cocaine exposure elicited excitatory responses in 48% of D1-MSNs and inhibitory responses in 72% of D2-MSNs (Figure 2E-F), a pattern markedly different from saline exposure (Figure S2E). In addition, D1-MSNs showed a significantly higher peak activity in response to cocaine than D2-MSNs (Figure 2G; Figure S2F-G). Next, to determine if D1- and D2-MSNs’ activity carries sufficient information to encode cocaine responses, we trained a support vector machine (SVM) decoder to distinguish baseline-vs cocaine-related activity (Figure 2H left). We show that the decoder accurately predicted cocaine exposure for D1- or D2-MSNs (91% and 87% accuracy, respectively; Figure 2H right). Importantly, when we trained a decoder to distinguish cocaine *versus* saline responses, we also show that both D1- and D2-MSNs differentially distinguish cocaine exposure from saline exposure (91% accuracy in both; Figure S2H-I).

Four weeks after cocaine exposure, mice were exposed to a single unexpected foot shock session, in which they received 10-foot shocks (0.7mA, 1s), delivered with an interval of 35-50s (Figure 2C). Individual neurons were classified as responsive based on activity changes 5s after shock (*p*<0.05, Permutation test). Foot shock predominantly excited both D1-MSN and D2-MSNs (81% and 82%, respectively; Figure 2I-J) with no significant differences in peak activity between populations (Figure 2K). Next, we trained an SVM to quantify how well D1- or D2-MSN population activity would predict foot shock exposure, i.e. distinguish activity pre- to post-foot shock delivery (Figure 2L left). We show that the decoder accurately predicted shock delivery for D1- or D2-MSNs (97% and 91% accuracy, respectively; Figure 2L right).

To assess how the same neuron responded to these opposing-valence stimuli, we analyzed the response of neurons that were tracked during cocaine and foot shock sessions. Importantly, tracked neurons’ activity was representative of the whole population activity (Figure 2M-N, 2P-Q). The majority of D1- and D2-MSNs were responsive to both stimuli (87% and 79%, respectively; Figure 2O, 2R). 38% of D1-MSNs responded in the same direction to both stimuli, in contrast with 8% of inhibited neurons. Regarding D2-MSNs, 14% of excited and 15% of inhibited neurons maintained the same type of response to both stimuli.

These findings reveal that cocaine and shock engage overlapping neuronal populations through distinct activity patterns: cocaine selectively excites D1-MSNs while inhibiting most of D2-MSNs, whereas shock broadly activates both populations.

### Optical reactivation of cocaine ensemble in the NAc elicits behavioral preference and positive reinforcement

Considering the differential anatomical distribution and functional properties of cocaine and foot shock ensembles, we next aimed to characterize the behavioral effects of optical re-activation of either ensemble. We used the TRAP2;Ai32 mouse model^29^, in which the presence of the 4-OHT-inducible Cre-recombinase (CreRT^T2^) enables permanent expression of channelrhodopsin (ChR2) in recruited cells (Figure 3A). After bilateral implantation of optic fibers into NAcC/mNAcSh (Figure S3A), mice were habituated to i.p. injections for 7 days, followed by saline or cocaine (20 mg/Kg) injections on TRAPing day plus i.p. administration of 4-OHT to induce expression of ChR2 (Figure 3B). Histological *pos-hoc* verification confirmed that cocaine administration recruited significantly more cells than saline in the NAcC and mNAcSh (Figure 3D-E).

**Figure 3.**
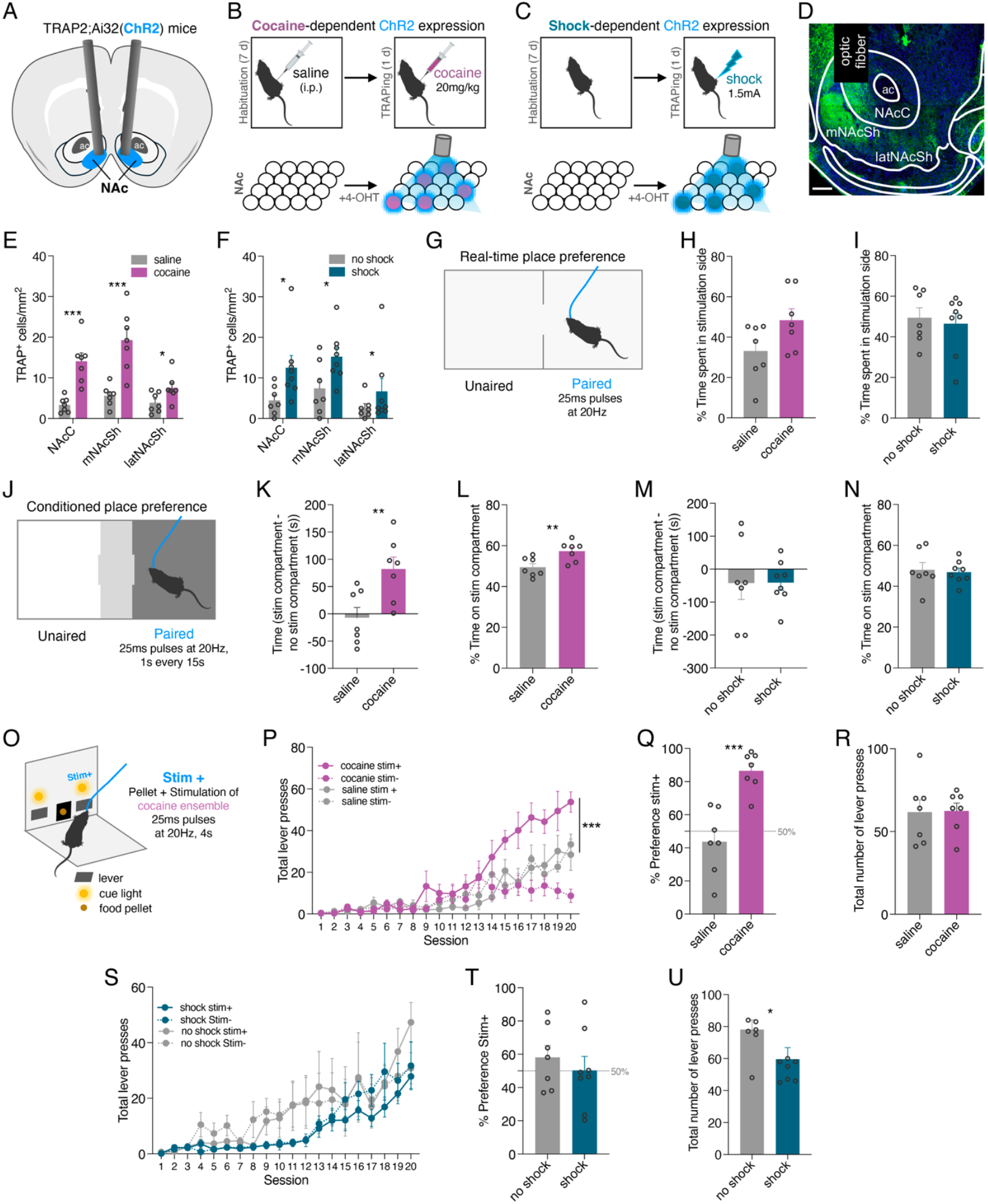
Optogenetic reactivation of cocaine-responsive ensembles, but not foot shock ensembles, in the NAc drives behavioural preference and reinforcement. **A** Schematic of the TRAP2;Ai32 mouse model used to tag and later reactivate cocaine- or shock-responsive neurons. Experimental design of **B** cocaine ensemble or **C** shock ensemble reactivation. **D** Representative expression of ChR2-eYFP, depicting optic fibber location; scale bar=250μm. Quantification of ChR2-expressing neurons after **E** cocaine administration (n_cocaine_=7, n_saline_=7; *unpaired t-test;* NAcC: *p*=0.0003; mNAcSh, *p*=0.0007; latNAcSh, *p*=0.0393) or **F** foot shock delivery (n_shock_=8, n_no shock_=7; NAcC, *Mann Whitney test*, U*p*=0.0103; mNAcSh, *unpaired t-test, p*=0.0336). **G** Schematic of the RTPP task. **H** Optical activation of cocaine ensembles produced a nonsignificant trend toward preference compared to saline controls (*unpaired t-test, p*=0.0742). **I** Optical activation of shock ensembles does not produce avoidance in comparison to no shock mice. **J** Schematic of the CPP task. Activation of cocaine ensembles induced significant CPP compared to saline ensemble stimulation, measured by **K** the difference in time spent between stimulation and non-stimulation compartments (*unpaired t-test, p*=0.0389) and by **L** percentage of time in the stimulation chamber (*p*=0.0093). **M, N** Activation of shock ensembles did not induce CPP in shock mice. **O** Schematic of the two-choice operant task. **P** Mice with cocaine ensemble stimulation developed a progressive preference for the stim+ lever over training days (*Mixed-effects two-way repeated-measures ANOVA*, group comparison, *p*=0.0021). On the last day, mice receiving optical excitation of cocaine ensembles presented **Q** a higher preference for the stim+ lever (*unpaired t-test, p*=0.0003), but **R** the same number of total lever presses. **R, S** Mice with shock ensemble stimulation did not develop preference for any lever but presented lower numbers of **T** total lever presses across on final training day. Data represent mean ± SEM. \**p*<0.05, \*\**p*<0.01, \*\*\**p*<0.001.

To evaluate the reinforcing properties of the cocaine ensemble, mice were first submitted to a real-time place preference (RTPP) task, in which optical stimulation was paired with one side of the apparatus (Figure 3G). Optical stimulation of cocaine ensemble was not sufficient to significantly induce behavioral preference (Figure 3H; Figure S3B), though a clear trend for preference for the stimulation side was observed. Next, we tested animals in the classical CPP test, in which one of the compartments was associated with optogenetic stimulation (Figure 3J). Cocaine ensemble reactivation induced a significant preference, measured by both the difference of time mice spent in the stimulation compartment in comparison with the time spent in the no-stimulation compartment (Figure 3K); as well as the percentage of time in the stimulation compartment (Figure 3L), in comparison with mice receiving optogenetic stimulation of saline ensemble. Notably, preference strength was positively correlated with the size of the cocaine ensemble in both NAcC and mNAcSh (Figure S3C), with no such correlation being found in the saline group, suggesting that larger ensembles encode stronger motivational value.

To further assess whether cocaine ensemble reactivation was sufficient to modulated reward-related behaviour, we tested mice in a two-choice lever pressing task, in which pressing either lever results in the delivery of a food pellet reward, with one lever being arbitrarily selected to deliver the pellet with simultaneous optogenetic stimulation of the cocaine (or saline) ensemble (stim+; Figure 3O). Throughout the session, mice can press either lever *ad libitum*; pressing one lever in one trial does not exclude the possibility of pressing the other lever in the next trial.

Mice receiving optical activation of cocaine ensemble showed a clear preference for stim+ lever in comparison with the stim-lever across acquisition days, significantly exceeding saline mice (Figure 3P-Q; no preference for either lever in saline group). No significant differences were found in the total number of lever presses (in both levers) in the last day of training between mice receiving stimulation of the cocaine ensemble and the saline ensemble (Figure 3R). When looking at the correlation between percentage for the stim+ lever and the size of the cocaine ensemble, we found a significantly positive correlation in the NAcC and mNAcSh (Figure S3D); no correlation was found in the saline group.

Together, these results indicate that activation of a cocaine ensemble in the NAc triggers behavioural preference and increases the value of a pellet reward, supporting a role in positive reinforcement.

### Reactivation of shock-responsive neurons of the NAc does not modify preference

Next, we tested whether optical re-activation of foot shock ensemble is sufficient to modulate behavioral preference. After the implantation of bilateral optic fibers into NAcC/mNAcSh (Figure S4A), mice were habituated to the foot shock apparatus and to i.p. injections for 7 days. On the following day, mice were exposed to a 10-minute session of unexpected foot shock delivery immediately followed by i.p. administration of 4-OHT (Figure 3C). Foot shock delivery recruited significantly more cells in comparison with no-shock group in the NAcC and mNAcSh (Figure 3F), confirming the presence of a shock ensemble.

In the RTPP, optical stimulation of the foot shock ensemble was not sufficient to induce behavioral preference/avoidance (Figure 3I; Figure S4B). Similarly, in the CPP test, optical activation of the foot shock ensemble also did not induce behavioral avoidance (Figure 3M-N). However, we found a negative correlation between the percentage of time spent in the stimulation compartment and the size of the foot shock ensemble in all NAc subregions (Figure S4C).

We next investigated the impact of re-activation of the foot shock ensemble in the two-choice task. Throughout acquisition days, mice receiving optical activation of the foot shock ensemble did not show preference or avoidance for either of the levers (Figure 3S-T), with learning curves resembling those of no-shock ensemble mice. Nonetheless, mice receiving optical stimulation of foot shock ensemble pressed significantly less in the stim+ than no-shock mice (Figure 3U), suggesting modest decreased motivated responses. Yet no correlation between preference for the stim+ lever and the size of the foot shock ensemble was found (Figure S4D).

These findings reveal an asymmetry in ensemble function: while cocaine ensembles drive positive reinforcement, shock ensemble does not produce optogenetic re-activation of cocaine or shock ensembles coupled with a recall of cocaine or shock would further amplify behavioral responses.

Saline- and cocaine-exposed mice were submitted to a cocaine CPP, where all animals received a cocaine injection (10 mg/Kg) before being conditioned to the compartments of the CPP; one of the compartments was additionally paired with optogenetic activation of the cocaine (or saline) ensemble (Figure 4A). Interestingly, while mice receiving optical activation of the cocaine ensemble only presented a trend for increased preference to spend more time in the stimulation chamber (Figure 4B-C), a clear positive correlation between preference for the stimulation side and size of the cocaine ensemble was observed in the NAcC and mNAcSh (Figure 4D; so correlation in saline group, Figure S5A). These findings suggest that reactivation robust avoidance, though it may dampen motivation to press for a lever associated with stimulation.

**Figure 4.**
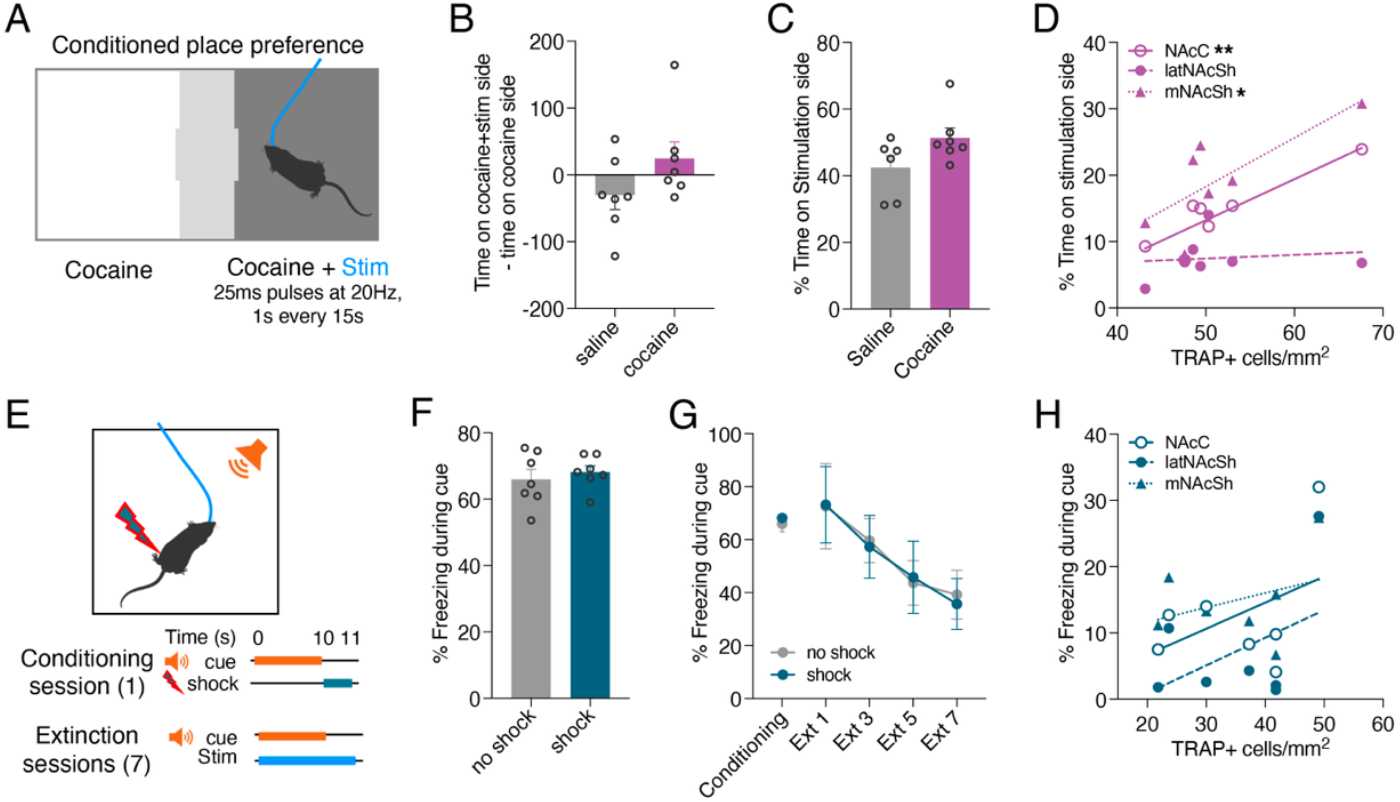
Recall of cocaine-(but not shock-) responsive neurons modulates behavioral preference. **A** Schematic of the cocaine CPP paradigm used to assess whether reactivation of cocaine- or saline-tagged ensembles during cocaine recall modulates preference; cocaine and saline mice received a cocaine injection (10mg/kg) before conditioning; one compartment was paired with optogenetic stimulation. Activation of cocaine ensembles did not alter cocaine CPP compared to saline ensemble stimulation, measured by **B** the difference in time spent between stimulation and non-stimulation compartments and **C** percentage of time spent in the stimulation chamber. **D** Correlation analysis between ensemble size and percentage of time spent in the stimulation chamber of the CPP test (positive correlations: *simple linear regression*; NAcC, *p*=0.0073; mNAcSh, *p*=0.0484). **E** Experimental design for the fear conditioning and extinction paradigm in mice with optogenetic stimulation of foot shock- or no-shock ensembles. **F** Freezing responses during fear conditioning. **G** Freezing responses across seven days of extinction. **H** Lack of correlation between freezing responses on the last extinction session and foot shock ensemble size. Data represent mean ± SEM. \**p*<0.05, \*\**p*<0.01.

### Recall of cocaine-but not shock-ensemble alters preference

Next, to determine whether ensemble reactivation modulates memory retrieval, we tested whether of cocaine-associated neuronal ensembles during drug recall enhances preference expression, supporting their role in reinforcing and amplifying cocaine-related motivational memory.

Next, we exposed mice previously exposed to foot shock or no-shock to a fear conditioning task. Thus, we trained mice to associate one 10-second auditory cue with the delivery of a mild foot shock (Figure 4E). Animals’ conditioned responses, measured as freezing behaviour, were similar between mice previously exposed to foot shock and no foot shock (Figure 4F). After the fear conditioning session, mice received 7 days of foot shock extinction, in which the foot shock-predicting cue and optogenetic stimulation were presented, but no foot shock was delivered. This approach intended to understand weather optical activation of the foot shock ensemble is sufficient to maintain fear responses or if mice would decrease conditioned responses due to extinction of the previous association. Mice of either group significantly decreased their freezing responses across the days of extinction (Figure 4G), suggesting that activation of the shock ensemble is not sufficient to maintain fear responses under extinction conditions. In accordance, no correlation between freezing responses on the last day of extinction and the size of the shock ensemble was found (Figure 4H; no correlation in no shock group, Figure S5B). These results suggest that reactivation of the foot shock ensemble does not preserve or reinstate fear behaviour, implying that additional circuit mechanisms are required to sustain conditioned fear responses.

## Discussion

This study demonstrates that within the NAc, rewarding and aversive stimuli leads to distinct patterns of cell-type recruitment and spatial distribution, and differential contribution for behaviour. Acute cocaine exposure recruits a D1-MSN-dominant ensemble localized to the anterior NAcC and latNAcSh, whereas unexpected foot shock engages an ensemble of D1- and D2-MSNs, broadly distributed across medial subregions. These findings align with previous studies showing that NAc cocaine-responsive ensembles are mainly composed of D1-MSNs^26,34^ and that these ensembles may reflect the divergent processing of stimuli with opposing valence within the NAc^35^.

The D1-MSN bias in cocaine ensemble aligns with established roles for dopaminergic signaling in reward processing. Cocaine-induced dopaminergic signalling promotes D1 receptor activation, facilitating synaptic potentiation, while D2 receptor activation decreases synaptic potentiation^24,34,36^. In contrast, foot shock engaged both D1- and D2-MSNs, suggesting a distributed representation of aversive information across accumbens pathways. This balanced recruitment may potentially occur through non-dopaminergic mechanisms and relying on dense glutamatergic inputs from other brain regions that largely encode aversive and fear responses such as the basolateral amygdala (BLA)^19,37–39^ or prefrontal cortex (PFC)^40^, to generate robust avoidance behavior. Moreover, the partial overlap observed between cocaine and shock ensembles indicates that a subset of accumbal neurons is capable of responding to stimuli of opposite valence, potentially reflecting shared circuits involved in salience or arousal rather than valence-specific encoding^16,41^.

Our calcium imaging data confirms and extends the anatomical findings, showing that D1- and D2-MSNs exhibit distinct yet complementary response profiles to cocaine and foot shock. Cocaine predominantly excited D1-MSNs while inhibiting D2-MSNs, consistent with previously reported data^42,43^, and supporting a dopamine-dependent gating of cocaine-dependent signals^7,44–47^. Conversely, both populations were robustly activated by foot shock. Importantly, the observation that many neurons responded to both stimuli but in an opposite direction supports the notion that the same neuronal populations can flexibly encode different valences depending on their inputs and neuromodulatory state, a phenomenon described for other brain regions^48–50^.

To establish a causal link between NAc ensemble activity and behavioural output, we optogenetically reactivated cocaine and shock ensembles. Re-activation of cocaine ensemble within the NAcC and mNAcSh was sufficient to elicit behavioral preference and positive reinforcement. In the CPP test, mice showed a clear bias toward the compartment paired with optogenetic stimulation of the cocaine ensemble, and this preference scaled with the size of the recruited ensemble. Similarly, in a two-choice operant task, stimulation of the cocaine ensemble reinforced lever pressing, indicating that the activity of these neurons carried intrinsic motivational value. These findings provide strong causal evidence that the ensemble of neurons activated during cocaine exposure encodes the positive valence and reinforcing properties of the drug, and that reactivation of this ensemble can recapitulate reward-seeking behavior. The correlation between ensemble size and behavioral strength further suggests that the extent of neuronal recruitment during drug experience may predict the intensity of subsequent motivational drive a concept that aligns with previous observations that larger ensembles underpin stronger sensitization to cocaine effects^51^.

Conversely, optogenetic reactivation of foot shock responsive ensembles failed to produce clear avoidance in either place preference or operant paradigms. While stimulation of the shock ensemble did not alter preference, it modestly decreased overall lever-pressing behavior, suggesting a general suppression of motivation rather than a directed aversive response. This dissociation implies that aversive ensembles in the NAc may primarily encode negative valence arising from its afferents from the BLA^50^, but lack the circuit-level output necessary to drive active avoidance when isolated from their broader network context. The absence of a strong behavioral phenotype upon shock ensemble reactivation may reflect the distributed nature of aversive processing across interconnected regions (reviewed in ^52^), which together orchestrate adaptive defensive behavior. Thus, while NAc ensembles contribute to the encoding of aversive experiences, their reactivation alone may not be sufficient to reproduce the full behavioral expression of avoidance, highlighting an asymmetry between reward- and punishment-related ensemble function. In line, when ensemble reactivation was combined with concurrent recall of the original experience, cocaine ensemble activation further enhanced conditioned preference, confirming its role in amplifying drug-associated motivational states^26,53^. Contrariwise, reactivation of the shock ensemble during fear extinction did not preserve freezing responses.

Together, these results highlight the NAc as a central hub for integrating and storing motivationally relevant information through valence-specific ensembles. Cocaine D1-MSN-enriched ensemble reactivation is sufficient to evoke positive reinforcement in the absence of external reward. Conversely, the limited behavioral influence of shock ensembles supports the idea that aversive processing in the NAc is more distributed and dependent on other outputs.

### Resource availability

#### Lead contact

Further information and requests for resources and reagents should be directed to and will be fulfilled by the lead contact, Carina Soares-Cunha.

## Data availability

Source Data are provided with this paper.

All data generated in this study will be deposited in the Zenodo database.

## Code availability

This paper reports original code that is available on the GitHub platform github.com/rewave-lab and published the identifier https://doi.org/10.5281/zenodo.13890127.

## Acknowledgments

This work was funded by the Portuguese Foundation for Science and Technology (FCT) under the scope of the project 2022.01467.PTDC (DOI: 10.54499/2022.01467.PTDC), a FEBS (Federation of European Biochemical Societies) Excellence Award and “Maria de Sousa” Award attributed to Carina Soares-Cunha by Bial Foundation and Órdem dos Médicos.

This work was also supported by the European Research Council (ERC) under the European Union’s Horizon 2020 research and innovation programme (grant agreement No 101003187) and by the “la Caixa” Foundation (ID 100010434), under the agreement LCF/PR/HR20/52400020. This work also received financial support from FEDER, COMPETE 2030 and national funds from FCT through the project COMPETE2030-FEDER-00708300 (DOI: 10.54499/2023.17631.ICDT) and FCT project 2024.14660.PEX.

CS-C and BC have Scientific Employment Stimulus contracts from FCT (2023.08896.CEECIND/CP2841/CT0023 (DOI: 10.54499/2023.08896.CEECIND/CP2841/CT0023); CEECIND/03898/2020 (DOI: 10.54499/2020.03898.CEECIND/CP1600/CT0015)). RC has an FCT PhD grant (2022.12973.BD; DOI: 10.54499/2021.06818.BD).

Host laboratory is funded by National funds, through FCT - projects UIDB/50026/2020 (DOI 10.54499/UIDB/50026/2020), UIDP/50026/2020 (DOI 10.54499/UIDP/50026/2020) and LA/P/0050/2020 (DOI 10.54499/LA/P/0050/2020).

## Author Contributions

CS-C conceived the project, designed the experiments and supervised the research; RC performed the behavioural experiments with TRAP;Ai14 and TRAP;Ai32 mice; AVD performed acquisition of calcium imaging data and histology; LAAA analysed most of the calcium imaging data; MM performed behavioural data analysis and histology; DVC performed RNAScope® and analysed the data; NVG assisted in behavioural performance; RMJ perform SVM from calcium imaging data; ET assisted in experiments with TRAP;Ai14 and TRAP;Ai32 mice; BC supported in TRAPing experiments with TRAP;Ai14 and TRAP;Ai32 mice; AJR supervised the research.

## STAR Methods

### Key resources table

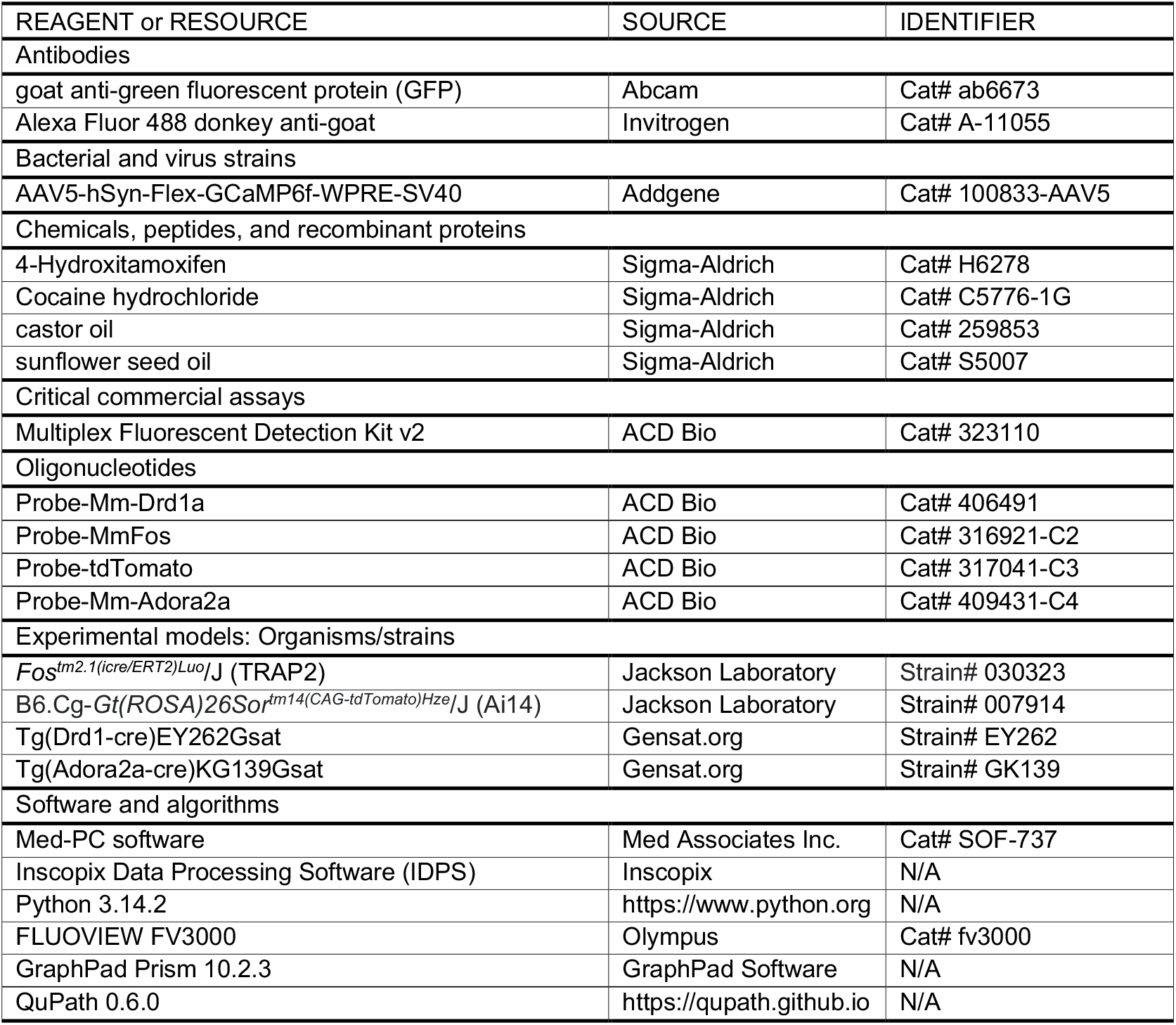

| REAGENT or RESOURCE | SOURCE | IDENTIFIER |
| --- | --- | --- |
| Antibodies |  |  |
| goat anti-green fluorescent protein (GFP) | Abcam | Cat# ab6673 |
| Alexa Fluor 488 donkey anti-goat | Invitrogen | Cat# A-11055 |
| Bacterial and virus strains |  |  |
| AAV5-hSyn-Flex-GCaMP6f-WPRE-SV40 | Addgene | Cat# 100833-AAV5 |
| Chemicals, peptides, and recombinant proteins |  |  |
| 4-Hydroxitamoxifen | Sigma-Aldrich | Cat# H6278 |
| Cocaine hydrochloride | Sigma-Aldrich | Cat# C5776-1G |
| castor oil | Sigma-Aldrich | Cat# 259853 |
| sunflower seed oil | Sigma-Aldrich | Cat# S5007 |
| Critical commercial assays |  |  |
| Multiplex Fluorescent Detection Kit v2 | ACD Bio | Cat# 323110 |
| Oligonucleotides |  |  |
| Probe-Mm-Drd1a | ACD Bio | Cat# 406491 |
| Probe-MmFos | ACD Bio | Cat# 316921-C2 |
| Probe-tdTomato | ACD Bio | Cat# 317041-C3 |
| Probe-Mm-Adora2a | ACD Bio | Cat# 409431-C4 |
| Experimental models: Organisms/strains |  |  |
| *Fos<sup>tm2.1(icre/ERT2)Luo/J</sup>* (TRAP2) | Jackson Laboratory | Strain# 030323 |
| *B6.Cg-Gt(ROSA)26Sor<sup>tm14(CAG-tdTomato)Hze/J</sup>* (Ai14) | Jackson Laboratory | Strain# 007914 |
| *Tg(Drd1-cre)EY262Gsat* | Gensat.org | Strain# EY262 |
| *Tg(Adora2a-cre)KG139Gsat* | Gensat.org | Strain# GK139 |
| Software and algorithms |  |  |
| Med-PC software | Med Associates Inc. | Cat# SOF-737 |
| Inscopix Data Processing Software (IDPS) | Inscopix | N/A |
| Python 3.14.2 | https://www.python.org | N/A |
| FLUOVIEW FV3000 | Olympus | Cat# fv3000 |
| GraphPad Prism 10.2.3 | GraphPad Software | N/A |
| QuPath 0.6.0 | https://qupath.github.io | N/A |

#### Experimental model and Subject details

Male and female heterozygous TRAP2;Ai14 and TRAP2;Ai32 mice (2-4 months old) were used. TRAP2;Ai14 and TRAP2;Ai32 mice were generated by crossing homozygous TRAP2 mice (#030323, Jackson Laboratories) with homozygous Ai14 mice (#007914, Jackson Laboratories), or Ai32 mice (#024109, Jackson Laboratories), respectively. For *in vivo* calcium imaging, male and female heterozygous D1-cre (line EY262, Gensat.org) and A2A-cre (line GK139, Gensat.org) transgenic mice lines (2-3 months old) with a C57BL/6J background were used. Animals were housed in groups of 3-5 and were maintained under standard laboratory conditions: 12h light/dark cycle (lights on at 8:00 am), temperature of 21 ± 1 ºC, and 60% relative humidity. Standard diet and water *ad libitum* were provided, except when stated otherwise. All behavioral experiments were performed during the light period of the light/dark cycle. Sample size used in behavioral tests was chosen according to previous studies; the investigator was not blind to the group allocation during behavioral performance, but it was blind in data analysis.

Health monitoring was performed according to FELASA guidelines. All animal experiments complied with the ARRIVE guidelines and were conducted according to European Union Regulations (Directive 2010/63/EU). Animal facilities and experimenters were certified by the Portuguese regulatory entity, Direção-Geral de Alimentação e Veterinária (DGAV). All protocols were approved by the Ethics Committee of the ICVS (ORBEA) and by DGAV (reference 008332).

### Method Details

#### Surgeries

Surgeries were performed under sterile conditions and sevoflurane (2–3%, plus oxygen at 1–1.5 l/min) anesthesia on a stereotaxic frame (Model 940, David Kopf Instruments, CA, USA). Throughout each surgery, mouse body temperature was maintained at 36°C using an animal temperature controller (ATC2000, World Precision Instruments, FL, USA). The mouse head was cleaned with 70% ethanol and a small incision from anterior to posterior was made on the skin to allow for aligning the head and drilling the hole for the injection site.

#### Imaging

Mice were unilaterally injected with 300nL of AAV5-hSyn-Flex-GCaMP6f-WPRE-SV40 (Addgene, MA, USA) into the NAc (coordinates^54^: +1.40 mm anteroposterior (AP), +0.7 mm mediolateral (ML), -4.3 mm dorsoventral (DV)) using a Nanojet III Injector (Drummond Scientific, PA, USA) at a rate of 1 nL/sec. The injection pipette was left in place for 10 minutes post-injection before it was removed. After the injection, a 0.6-mm-diameter gradient index (GRIN) lens with a baseplate attached (Inscopix, Inc., CA, USA) was slowly lowered into the NAc (0.2 mm/min) directly above the injection site after a slow pre-track was made with a 26-gauge blunt needle (until 0.4 mm above the target DV coordinate). Once in place, the craniotomy was closed with a low-toxicity silicone adhesive (Kwik-Sil, World Precision Instruments, FL, USA) and the lens was secured to the skull using dental cement (Superbond C&B kit, Sun Medical, Shiga, Japan).

#### Optogenetics

TRAP2;Ai32 mice were bilaterally implanted in the NAc (coordinates^54^: +1.4 mm AP, ±0.9 mm ML, and -4.3 mm DV) with optical fibers with 1.25 mm stainless steel ferrule (200 μm core; Thorlabs, NJ, USA), secured to the skull with dental cement (C&B kit, Sun Medical, Shiga, Japan).

After surgery, mice were removed from the stereotaxic frame and were allowed to recover from the anesthesia in their home cage under a heating lamp. All animals were treated 30 minutes before surgery with an analgesic – 0.1 mg/Kg buprenorphine (i.p. injection; Bupaq, Richter Pharma, Wels, Austria) and maintained for 48h after surgery. For calcium imaging, mice were left to recover for six weeks, while animals used for optogenetics were left to recover for two weeks before starting behavioral procedures.

#### Optogenetic manipulation

Optogenetic stimulation was performed using 5 mW blue light (at the tip of the fiberoptic) generated by a 473nm DPSS laser (CNI Laser, Changchun, China) and a pulse generator (Master-8; AMPI, MN, USA), delivered through fiberoptic patch cords (0.22 NA, 200 mm diameter; Thorlabs, NJ, USA) attached to the implanted ferrules.

The stimulation was performed as follows: 25 ms light pulses at 20 Hz, 50% duty cycle. The duration of the stimulation varied with the behavioral protocol.

#### Drug preparation

4-hydroxytamoxifen (4-OHT, H6278, Sigma-Aldrich, Merck KGaA, Darmstadt, Germany) was dissolved at a concentration of 50 mg/mL in 100% ethanol followed by a solution of 1:4 castor oil:sunflower seed oil (259853 and S5007, Sigma-Aldrich, Merck KGaA, Darmstadt, Germany). The final concentration was 5 mg/mL, and all mice were injected intraperitoneally (i.p.) at a dose of 50 mg/Kg.

2 mg/mL of cocaine hydrochloride (C5776-1G, Sigma-Aldrich, USA) was prepared by dissolving the powder in 0.9% saline solution. Mice were administered i.p. at a dose of 20 mg/Kg.

### Behavioral experiments

#### Ensemble Labeling – TRAPing

No Shock and Shock mice were habituated to the foot shock apparatus (21.59cm x 18.08cm; Med Associates Inc., VT, USA; 10 min; free exploration) and all animals (No Shock, Shock, Saline, and Cocaine) were habituated to i.p. injections (200 μl saline; 25G x 5/8 needle; administered in the chemical hood) for 7 days. After each session of habituation, mice were individually housed for 6h, returning to their homecages after this period.

On the TRAPing day, mice underwent a 10-minute foot shock task where Shock mice received 20 1.5mA foot shocks generated by a shocker module (Med Associates Inc., VT, USA), delivered every 30 seconds. No Shock mice were placed in the apparatus for 10 minutes, and the houselight turned on, but no foot shock was delivered. A computer equipped with Med-PC software (Med Associates Inc., VT, USA) was used to control the equipment. Cocaine and Saline mice received an injection of cocaine (20 mg/Kg, i.p.) or saline, respectively. After each session, all groups received a 4-OHT injection to allow labeling of the cells that responded to the specific stimuli. Mice were individually housed for 6h to prevent the neurons from being activated by external stimuli (i.e., social interaction with other mice) and returned to their homecages after the isolation period.

#### Ensemble Labeling - endogenous c-fos expression

Two weeks after the TRAPing day, and after 7 days of habituations, TRAP2;Ai14 mice were exposed to the opposite stimulus and were euthanized 60 minutes after the end of the session, to allow the detection of endogenous c-fos protein induced by the stimulus exposure.

#### Real-time place preference (RTPP)

The RTPP test was adapted from a previously described protocol^55^. Briefly, a habituation session of 15 min was performed, in which mice were allowed to freely explore the apparatus (54cm x 48cm, divided in half), with the patch cables attached to the animal’s optic ferrules. On the next day, a 20-minute session was performed, in which one of the sides of the apparatus was designated as the stimulation-paired side. Mice started the behavioral performance at the center of the apparatus, and optical stimulation was applied during the entire period that the animal stayed on the stimulation-paired side. The attribution of the side paired with optical stimulation was done randomly. Behavior was recorded via a video camera and time spent in each chamber was manually evaluated, blindly. Data is presented as the percentage of time spent on the stimulation side.

#### Conditioned place preference (CPP)

The CPP test was based on previously described protocols^55,56^. Mice were first exposed to a 15-min pre-test session, followed by 6 conditioning sessions (30 min; 3 conditionings to the stimulation-paired chamber, and 3 conditionings to the no-stimulation chamber, altered between conditions), and a 15-min post-test session. In both pre-test and post-test sessions, mice were placed in the middle compartment and freely explored the apparatus (two chambers – 30cm x 21cm – divided by a middle compartment – 12cm x 21cm; Med Associates Inc., VT, USA). In the conditioning sessions mice were confined to one of the chambers that was randomly signed as the one associated with or without optical stimulation. In the CPP with cocaine (20 mg/Kg), mice received an i.p. injection before the beginning of each conditioning session. Optical stimulation consisted of 1-sec every 15 seconds. A computer equipped with Med-PC software (Med Associates Inc., VT, USA) was used to control the equipment and record data, allowing assessment of time spent in each chamber. Data is presented as the percentage of time spent on the stimulation-paired chamber during the post-test session and the difference of time spent on the stimulation-paired chamber between the post-test and the pre-test sessions.

#### Two-choice schedule of reinforcement task

Mice were placed and maintained on food restriction (≈ 2 g/day of standard laboratory chow) to maintain 80-90% free-feeding weight. Behavioral sessions were performed in operant chambers (Med Associates Inc., VT, USA) containing a central magazine that provided access to 20 mg standard food pellets (F0021, Dustless Precision Pellets, Bio-Serv, NJ, USA), two retractable levers located on each side of the magazine with cue lights above them, and a 2.8 W, 100 mA house light positioned at the top center of the wall opposite to the magazine to provide illumination.

Mice were trained for a food-seeking operant task with optogenetic stimulation during 20-minute daily sessions^57^. A single press on the stim+ lever led to the delivery of one 20 mg food pellet plus optical stimulation (25 ms pulses delivered at 20 Hz over 4 s) paired with a 4-second auditory cue. Single press on the reward alone (stim–) lever led to the delivery of one food pellet paired with another 4-second auditory cue, but with no laser stimulation. For both levers, presses during the 4-s after pellet delivery have no further consequence. Each daily session begins with a single lever presented alone to allow the opportunity to earn its associated reward (either stim+ or stim−), after which the lever is retracted. Then, the alternative lever is presented by itself to allow the opportunity to earn the other reward. Finally, both levers are extended together for the remainder of the session. A computer equipped with Med-PC software (Med Associates Inc., VT, USA) controlled the equipment and recorded the data. Data is presented as the number of lever presses for each lever, and the percentage of preference for the stim+ lever.

#### Fear conditioning

No Shock and Shock animals underwent a fear conditioning task. All sessions started with 60 s of habituation period, with the house light on. The conditioned stimulus (CS) consisted of an 80 dB, 2-kHz tone plus a cue light, paired with a mild foot shock (US; 0.5 mA, over 1 s). During the conditioning session, mice were exposed to 10 CS-US pairings. Each trial consisted of an 11-s tone, that co-terminated with the foot shock delivered through the stainless-steel grid floor, followed by a random intertrial interval (ITI) of 25-40 seconds. During the next 7 days, mice underwent extinction sessions, in which the CS was presented for 11s, coupled with optical stimulation, but no foot shock was delivered. All sessions were recorded via a video camera. Data is presented as the percentage of time in which mice freeze during the CS period. The freezing response was defined as the time that mice spent immobile (lack of any movement including sniffing, except respiration) during the CS period and calculated as percentage of total cue time ((freezing time/cue duration) x 100).

#### Calcium Imaging Acquisition

GCaMP6f fluorescence signals were recorded using a miniature integrated fluorescence miniscope (Inscopix, Palo Alto, CA) through GRIN lenses implanted in the NAc of freely behaving mice. Before each imaging session, the microscope was carefully attached to the baseplate while the mouse was gently restrained. The analog gain (4-6), LED output power (0.8-1.5mW) and focal plane were kept constant for each subject across all imaging sessions.

To synchronize foot shock events with imaging data, the nVoke Imaging System’s Data Acquisition Box (Inscopix) was triggered by the pyControl behavioral software. To synchronize i.p. injection events with imaging data, a synchronized TTL signal was inputted into the Imaging System’s Data Acquisition Box (Inscopix). Compressed grayscale images were acquired at 20 frames per second with a spatial down sampling factor of four.

Calcium imaging was conducted during cocaine, saline and foot shock sessions. The number of neurons recorded per animal is presented in Supplementary Table 1.

#### Calcium Imaging Data Processing

##### Data Preprocessing

Calcium imaging data were preprocessed using the Inscopix Data Processing Software (IDPS). The field of view was first cropped to remove peripheral regions, and this same area was maintained across all recordings for each animal. A low-pass filter was then applied to reduce noise, followed by an initial motion correction using standard IDPS settings. The resulting data were exported as a single TIFF image stack. A second motion correction was subsequently carried out in Python using the CaImAn toolbox^58^, which implements the NoRMCorre algorithm. This non-rigid motion correction method efficiently reduced motion-related artifacts by optimizing local spatial transformations across frames.

After motion correction, source extraction was performed using the extended constrained non-negative matrix factorization algorithm for one-photon imaging (CNMF-E^59^) implemented in CaImAn. This procedure automatically identified regions of interest (ROIs) and produced denoised and deconvolved fluorescence traces (F). To normalize these signals, a custom Python script was used to estimate the baseline fluorescence (F_0_) via CaImAn’s internal function GetSn, yielding normalized fluorescence activity (ΔF/F_0_). All automatically identified ROIs were manually inspected to remove irregularly shaped or noisy components and to confirm that each ROI corresponded to a neuron.

##### Cell Registration

Following manual curation, the spatial footprints of confirmed neurons were extracted from the CNMF-E output for each session. To track cells across sessions, two complementary registration methods were used. The first method employed CellReg^60^, a MATLAB-based package that aligns spatial footprints across sessions through translation and rotation, using the first session as the reference map. For each cell pair, CellReg computes a probability (P_same) that the two footprints belong to the same neuron, based on spatial correlation and centroid distance. Cells were considered matched if P_same exceeded 0.5.

The second approach used the internal registration function of CaImAn, register_multisession, which compares spatial footprints across sessions using intersection-over-union metrics and resolves cell correspondences using the Hungarian algorithm. The first session was again used as the reference. To allow greater flexibility than CellReg, the “maximum distance considered” was set to 0.9 and the “maximum centroid distance” to 100.

The results obtained from both registration methods were combined and verified through manual inspection to ensure consistent and reliable cell tracking across all imaging sessions.

#### Calcium Data Analysis

##### Alignment of Activity to Behavioral Events

For cocaine or saline exposure, i.p. events were recorded via an external TTL signal triggered manually, and activity was analyzed within 300-second pre- and post-TTL windows.

For foot shock exposure, neuronal activity was aligned to the onset of the shock, with 5-second pre- and post-shock windows analyzed.

##### Permutation Test

To determine neuronal responses to stimuli, changes in the average normalized fluorescence signal (ΔF/F_0_) were evaluated using a permutation test. For each cell, the fluorescence trace within a 300-second window for cocaine or saline or a 5-second window for foot shock was randomly shuffled 1000 times. If the absolute difference between the real pre- and post-stimulus averages exceeded 95% of the differences observed in the shuffled data, the neuron was classified as responsive. Neurons showing an increase in post-stimulus activity were classified as excited, whereas those showing a decrease were classified as inhibited. Cells without significant changes were classified as non-responsive.

##### Signal Normalization (z-score)

For subsequent analyses, responses to stimuli were expressed as z-scores to quantify the relative intensity of neuronal activation. The z-score was calculated as:

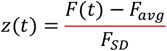

where *F(t)* represents the normalized fluorescence signal (ΔF/F_0_), and *F_avg* and *F_SD* are the mean and standard deviation of ΔF/F_0_ calculated across all pre-stimulus baseline periods for all trials.

##### Heatmaps

Heatmaps of neuronal activity were generated using the seaborn package in Python (seaborn.heatmap), based on the mean z-scores of individual cells. The plotted interval ranged from -3 to +3 seconds for foot shock and from -300 to +300 seconds for cocaine or saline, with time zero corresponding to event onset. In all heatmaps, the color bar was set with the “extend” option enabled to represent data beyond the upper and lower bounds of the scale.

##### Peak analysis

The maximum peak was calculated for each group during the period following stimulus onset using the Python package scipy.find_peaks function (thresholds: minimum prominence = 2.5; minimum peak height = 1.96).

##### Population Decoder Analysis

For population-level decoding, we constructed population vectors which size corresponded to the number of cells. Each element of a vector represented the mean z-score of a single cell either during the stimulus-evoked response window or during the preceding baseline. In datasets containing multiple stimulus presentations, each trial provided a baseline–response pair of population vectors. In datasets with only a single prolonged stimulus per recording, trials were instead defined by dividing the baseline and post-event periods into non-overlapping time windows, and population vectors were computed for each segment. In all sessions, this procedure yielded paired baseline–response vectors characterizing population activity around the stimulus.

The linear support vector machine (SVM) classifier was trained on 70% of the population vectors, with the remaining 30% being reserved for testing decoding performance. In within-recording analyses, the SVM was trained to discriminate response vectors from their corresponding baselines to assess whether population activity carried stimulus-related information. In cross-recording analyses, SVMs were trained to classify response vectors from different stimuli to test whether population responses were separable across stimulus identities. For both within- and cross-recording analyses, we generated 1000 independent SVM models for the real and shuffled datasets, and decoding accuracy was defined as the mean performance across these models.

##### Sacrifice and brain sectioning

Mice were deeply anesthetized and transcardially perfused with 0.9% saline, followed by 4% paraformaldehyde (PFA). For immunohistochemistry, whole heads with the optic fibers attached were immersed for 24 h in 4% PFA so that the track is clearly visible for histological analysis. Then, brains were removed, rinsed and stored in 30% sucrose at 4 °C until sectioning. Coronal sections (40 μm) were performed on a vibrating microtome (VT1000S, Leica, Germany) and slices were stored at 4 °C on 12-well plates (or long-term storage in cryoprotectant solution at −20 °C) until use. Slices from the area of interest were selected using the Mouse Brain Atlas^54^. For RNAscope *in situ* hybridization, brains were removed, post-fixed in 4% PFA for 24 hours at 4 ºC under agitation and subsequently transferred to sucrose solutions of increasing concentrations (10%, 20%, and 30%), progressing to the next concentration once the brains had sunk to the bottom of the container. Brains were frozen in Optimal Cutting Temperature (OCT) embedding medium (Tissue-Tek, Sakura Finetek USA, Inc., CA, USA) with isopentane on dry ice and stored at –80 ºC until sectioning. Sections (16 mm) were cut using a cryostat (Leica CM1900; Leica Biosystems, Nussloch, Germany).

##### Immunohistochemistry

Brain slices were washed with phosphate-buffered saline (PBS), permeabilized with PBS containing 0.3% Triton X-100 (PBS-T 0.3%), blocked with PBS-T plus 10% fetal bovine serum (FBS; Invitrogen, MA, USA), and incubated with the primary antibody goat anti-green fluorescent protein (dilution 1:500; ab6673, Abcam, Cambridge, UK) overnight at 4ºC under agitation. After washing with PBS-T, sections were incubated with the secondary fluorescent antibody Alexa Fluor 488 donkey anti-goat (dilution 1:500; A-11055, Invitrogen, CA, USA). All antibodies were diluted in PBS-T with 2% FBS. Slices were washed with PBS-T, incubated with 4’,6-diamidino-2-phenylindole (DAPI, 1:1000; 62248, Thermo Scientific™, MA, USA) for nucleus staining, washed with PBS and mounted using Immu-Mount™ Mounting Medium (Epredia™, 9990402, NH, USA). Slides were stored at 4 °C and kept protected from light.

##### RNAscope® *in situ* hybridization

Brain slices were used to visualize and amplify target RNA: D1-MSN (Probe-Mm-Drd1a, Cat #406491), c-fos (Probe-MmFos, Cat #316921-C2), tdTomato (Probe-tdTomato, Cat #317041-C3), and D2-MSN (Probe-Mm-Adora2a, Cat #409431-C4) using the RNAscope® *in situ* hybridization Multiplex Fluorescent Detection Kit v2 (Cat #323110, Advanced Cell Diagnostics, ACD, Bio-Techne, CA, USA), following the manufacturer’s instructions. Briefly, slides were post-fixed, dehydrated, and incubated in hydrogen peroxide as instructed. Tissue pretreatment was performed by steaming with target antigen retrieval reagent (Cat #322000, ACD, Bio-Techne, CA, USA) for 30 min. Protease III (Cat #322337, ACD, Bio-Techne, CA, USA) was applied to the sections and incubated at 40 ºC for 30 min, followed by hybridization with the selected probes at 40 ºC for 2 h. Signal amplification was performed using the provided AMP1-4 detection reagents, applied sequentially, and incubated for 15–30 min each at 40 ºC, with washing steps in 1X RNA scope between each reagent. The C4 channel probe was developed with the RNAscope 4-Plex Ancillary Kit (Cat #323120, ACD, Bio-Techne, CA, USA). Slices were then incubated with fluorophore dyes (Opal 520 (Cat #FP1487001KT), Opal 570 (Cat #FP1488001KT), Opal 620 (Cat #FP1495001KT), and Opal 690 (Cat #FP1497001KT), Akoya Biosciences Inc., MA, USA), with each channel developed using its corresponding HRP signal. Tissue was counterstained with DAPI, and slides were mounted in Immu-Mount™ Mounting Medium (Epredia™, 9990402, NH, USA), left to dry overnight in the dark, and stored at 4ºC, until image acquisition.

##### Image acquisition and analysis

For immunohistochemistry, images were collected and analyzed by confocal microscopy (FLUOVIEW FV3000, Olympus). Quantification of ChR2 positive cells was performed with ImageJ version 1.42 software^61^. For RNAscope® *in situ* hybridization, images were acquired at 20X on a Phenoimager™ HT (Akoya Biosciences Inc., MA, USA) and quantifications were done semi-automatically using QuPath software. NAc slices were overlaid with the mouse brain atlas^54^ to estimate the stereotaxic coordinates. *tdTomato, Drd1* and *A2a* fluorescent images were overlaid to identify co-localization of mRNA expression, defined by the presence of at least ten distinct fluorescent puncta. Co-localization of *tdTomato* and *C-fos* mRNA expression was also analyzed.

##### Statistical analysis

The presence of outliers was tested using ROUT (Q = 1%) test; outliers were removed before advancing in the data analysis. Normality tests (K-S) were performed, and statistical analysis was conducted accordingly. Student’s t-test (normality assumed) or Mann-Whitney (normality not assumed) tests were performed for comparisons between the two groups. Ordinary Two-Way Analysis of Variance (ANOVA; normality assumed) or Friedman (normality not assumed) tests were used for comparisons between groups. Repeated measures two-way ANOVA test was performed for comparisons between groups across days. Proper post-hoc multiple comparisons tests were used for group difference determination (Sidak test). Results are presented as mean ± standard error of the mean (SEM) and were considered statistically significant for *p* < 0.05. Statistical details of experiments can be found in figure legends. The final number of animals is depicted in figure legends. All statistical analyses were performed using GraphPad Prism 10.2.3 (GraphPad Software, Inc., La Jolla, CA, USA).

## Supplemental data

**Figure S1.**
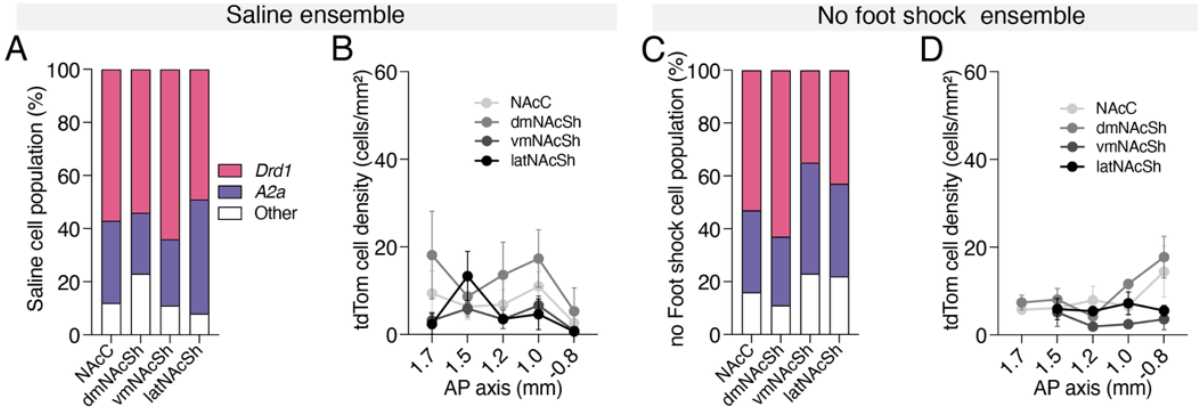
Neuronal ensemble recruitment in the NAc following saline or foot shock exposure. **A** tdTom+ cell-type composition within NAc sub-regions of no saline ensembles. **B** The saline ensemble displayed few labelled cells and a less defined anteroposterior distribution than the cocaine ensemble. **C** tdTom+ cell-type composition within NAc sub-regions of no foot shock ensembles. **D** The no shock ensemble displayed few labelled cells than the shock ensemble and a generalized anteroposterior distribution. Data represent mean ± SEM.

**Figure S2.**
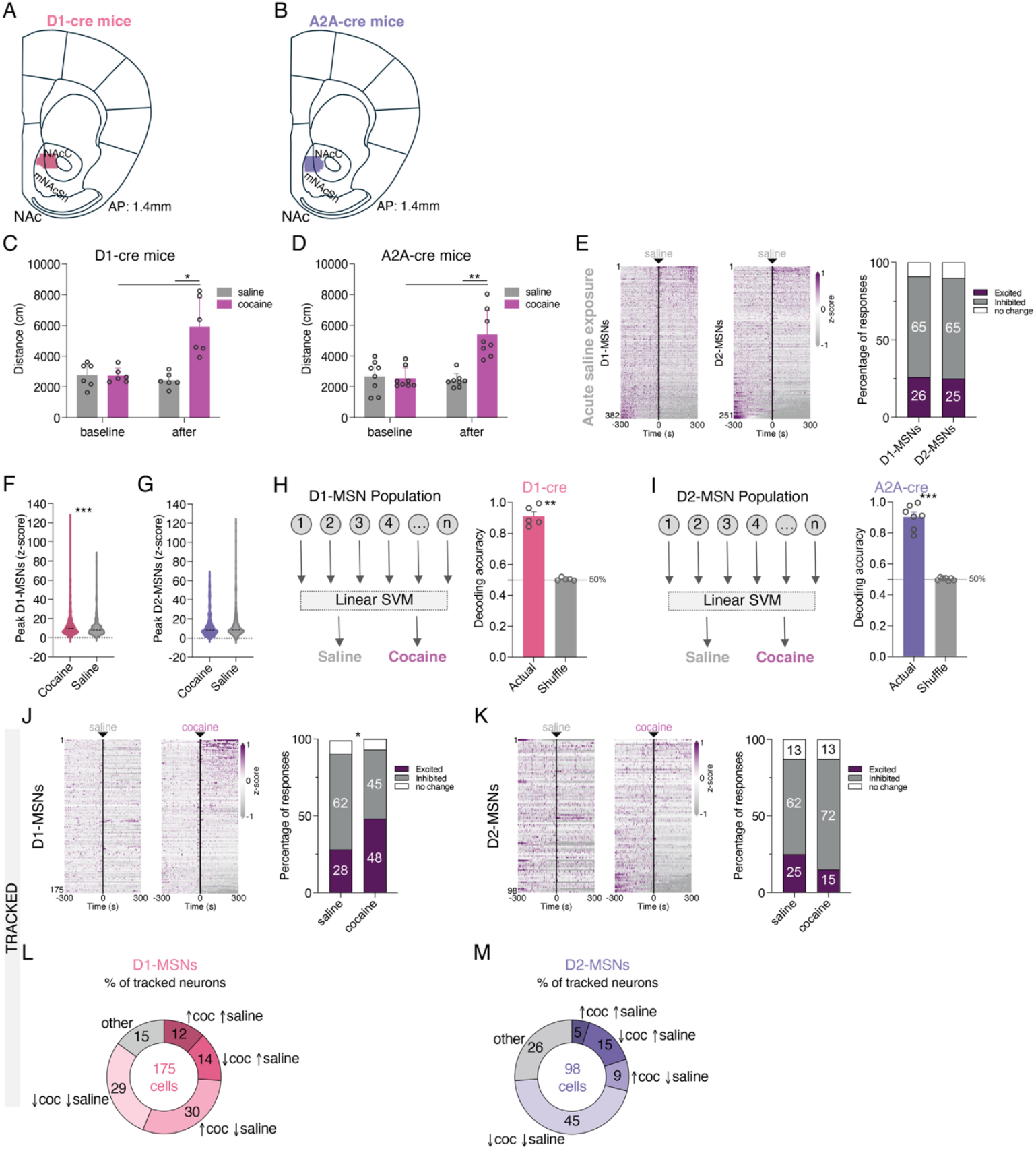
GRIN lenses placement of D1- and A2a-cre mice and D1- and D2-MSN calcium dynamics during saline and cocaine exposure. Anatomical localization of GRIN lens placement the NAc of **A** D1-Cre and **B** A2A-Cre mice; core to medial shell; anteroposterior (AP): 1.4mm. **C** Locomotor activity before and after cocaine or saline injection in D1-cre mice, showing that cocaine administration caused hyperlocomotion (*2-Way ANOVA*; Group comparison: F_1,5_=24.9, *p*=0.0041; *Sidak’s test*, baseline cocaine *versus* after cocaine *p*=0.0289, after saline *versus* after cocaine *p*=0.0126). **D** Locomotor activity before and after cocaine or saline injection in A2A-cre mice, showing that cocaine administration caused hyperlocomotion (*2-Way ANOVA*; Group comparison: F_1,7_=16.6, *p*=0.0047; *Sidak’s test*, baseline cocaine *versus* after cocaine *p*=0.0066, after saline *versus* after cocaine *p*=0.0050). **E** Activity heatmaps of individual D1-(n=382 cells) and D2-MSNs (n=251 cells) aligned to saline administration (left) and percentage of excited, inhibited and non-responsive D1- and D2-MSNs (right). **F** Peak z-score quantification of individual D1-MSNs after cocaine or saline administration (*Mann Whitney test*, U=69242, *p*=0.0003). **G** Peak z-score quantification of individual D2-MSNs after cocaine or saline administration. **H** SVM decoder accuracy for predicting cocaine trials from saline trials, based on D1-MSN population activity (*Mann-Whitney*, U_8_=0, *p*=0.0079). **I** SVM decoder accuracy for predicting cocaine trials from saline trials, based on D2-MSN population activity (*Mann-Whitney*, U_12_=0, *p*=0.0006). **J** Activity heatmaps of individual D1-MSNs (n=175 cells) tracked over the cocaine or saline administration sessions (left), and percentage of excited, inhibited and non-responsive D1-MSNs in response to cocaine or saline administration (right) showing higher proportion of excited neurons after cocaine administration (*xi-square test, p*=0.0148). **K** Activity heatmaps of individual D2-MSNs (n=98 cells) tracked over the cocaine or saline administration sessions (left), and percentage of excited, inhibited and non-responsive D2-MSNs in response to cocaine or saline administration (right). Percentage of each type of response of neurons tracked during the cocaine session and saline session for **L** D1-MSNs and **M** D2-MSNs. Data represent mean ± SEM. \**p*<0.05, \*\**p*<0.01, \*\*\**p*<0.001.

**Figure S3.**
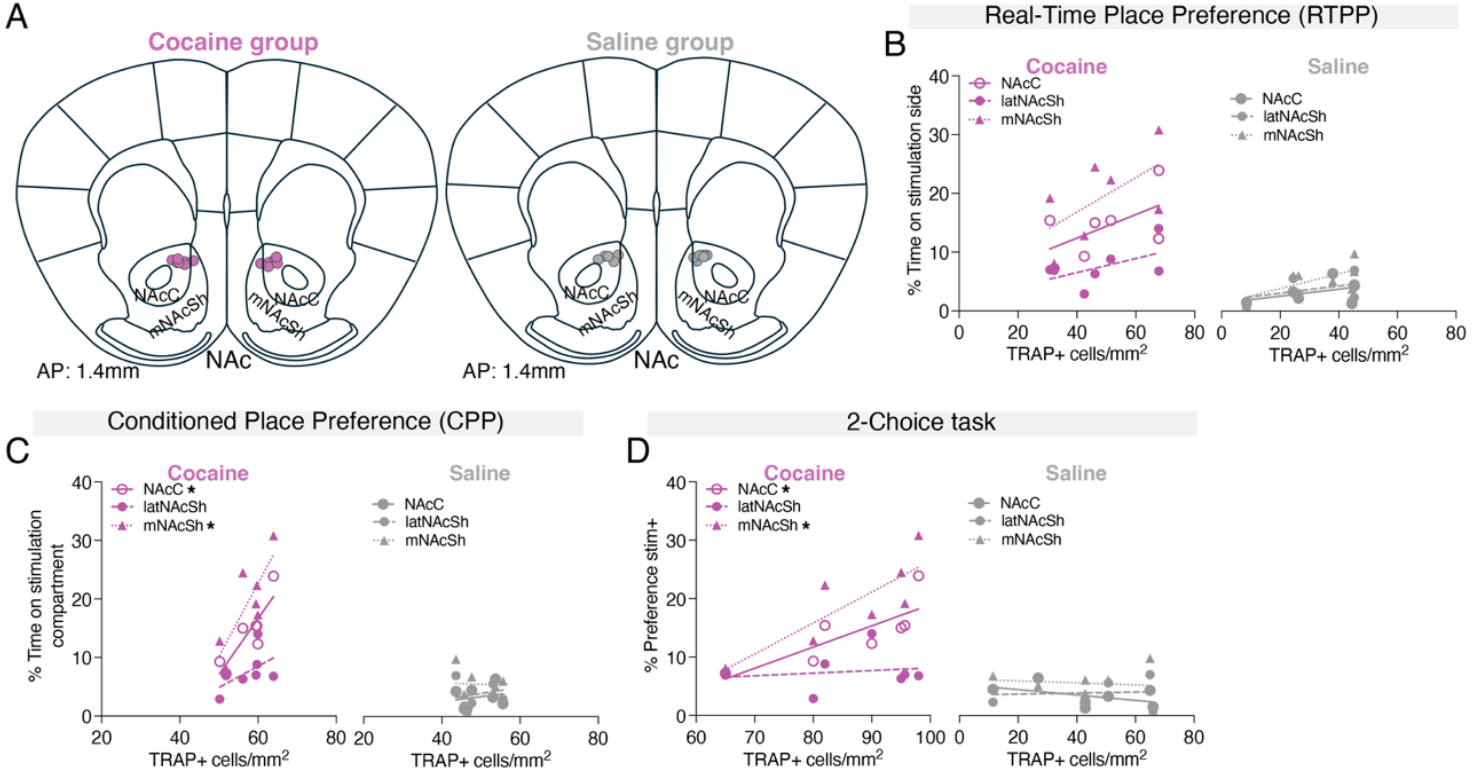
Anatomical location of optic fibbers in cocaine and saline groups and correlation analysis between behavioural performance and number of TRAPed cells. **A** Schematic illustration of optic fibber placements targeting the NAcC and mNAcSh in TRAP2;Ai32 mice used for optogenetic activation of cocaine or saline ensembles (AP: 1.4mm). Correlation analyses between ensemble size and behavioural preference of cocaine (left) or saline (right) groups in the **B** RTPP, **C** CPP (positive correlations: *simple linear regression*; NAcC, r^2^=0.7522, F_1,5_=15.2, *p*=0.0115; mNAcSh, r^2^=0.6596, F_1,5_=9.7, *p*=0.0265), and **D** 2-choice (positive correlations: *simple linear regression*; NAcC, r^2^=0.6249, F_1,5_=8.3, *p*=0.0343; mNAcSh, r^2^=0.6840, F_1,5_=10.8, *p*=0.0217) tests. \**p*<0.05.

**Figure S4.**
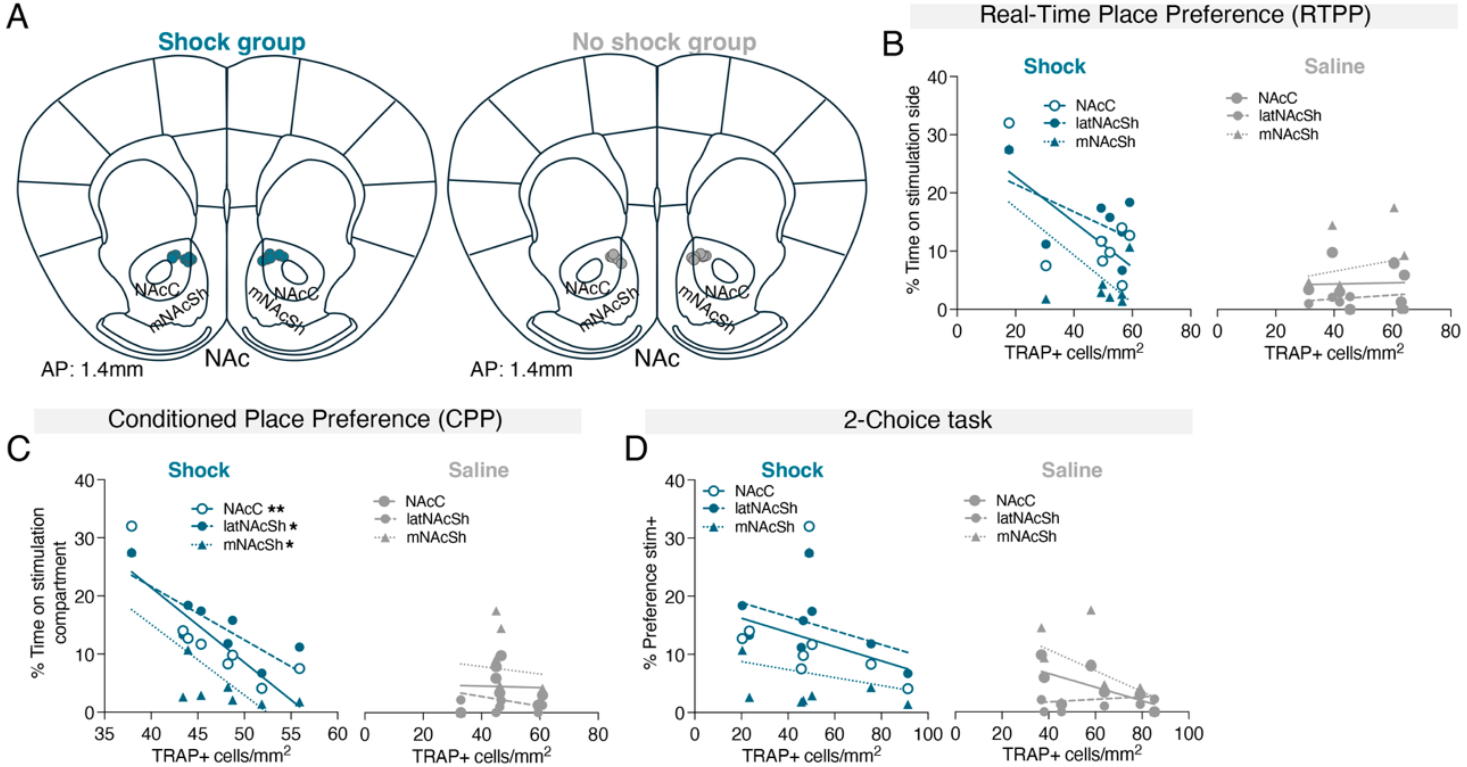
Anatomical location of optic fibbers in shock and no shock groups and correlation analysis between behavioural performance and number of TRAPed cells. **A** Schematic illustration of optic fibber placements targeting the NAcC and mNAcSh in TRAP2;Ai32 mice used for optogenetic activation of shock or no shock ensembles (AP: 1.4mm). Correlation analyses between ensemble size and behavioural preference of shock (left) or no shock (right) groups in the **B** RTPP, **C** CPP (positive correlations: *simple linear regression*; NAcC, r^2^=0.7096, F_1,6_=14.7, *p*=0.0087; mNAcSh, r^2^=0.5655, F_1,6_=7.8, *p*=0.0314; latNAcSh, r^2^=0.6786, F_1,7_=12.7, *p*=0.0119), and **D** 2-choice tests. \**p*<0.05, \*\**p*<0.01.

**Figure S5.**
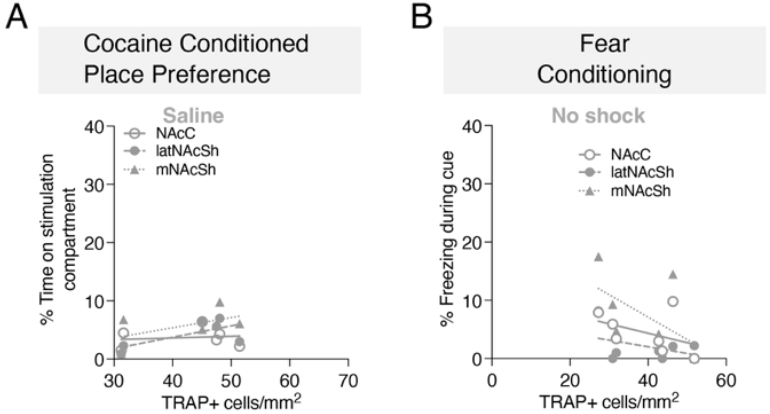
Correlation analysis between behavioural performance and number of TRAPed cells in saline and no shock groups during recall. **A** Correlation analysis between ensemble size and behavioural preference of saline group in the cocaine CPP. **B** Correlation analysis between ensemble size and behavioural preference of no shock group in the last extinction session of the fear conditioning task.

**Table S1.** Number of neurons recorded per animal in each behavioral session.

|  | Number of Recorded neurons in session |  |  |  |  |
| --- | --- | --- | --- | --- | --- |
|  | Saline | Cocaine | Foot Shock | Tracked sessions |  |
|  |  |  |  | Saline/Cocaine | Cocaine/Foot Shock |
| D1-cre mice |  |  |  |  |  |
| Animal 1 | 3 | 5 | 9 | --- | --- |
| Animal 2 | 108 | 120 | 89 | 48 | 62 |
| Animal 3 | 129 | 148 | 125 | 71 | 103 |
| Animal 4 | 65 | 67 | 85 | 34 | 44 |
| Animal 5 | 52 | 41 | 50 | 17 | 21 |
| Animal 6 | 25 | 45 | 14 | 5 | 7 |
| Total D1-MSNs | 382 | 426 | 372 | 175 | 237 |
| A2A-cre mice |  |  |  |  |  |
| Animal 7 | 14 | 22 | 14 | 8 | 8 |
| Animal 8 | 3 | 2 | 14 | --- | --- |
| Animal 9 | 40 | 32 | 27 | 17 | 18 |
| Animal 10 | 22 | 18 | 18 | 6 | 6 |
| Animal 11 | 44 | 46 | 51 | 25 | 32 |
| Animal 12 | 73 | 53 | 99 | 39 | 45 |
| Animal 13 | 8 | 9 | 1 | 1 | --- |
| Animal 14 | 47 | 27 | 15 | 2 | --- |
| Total D2-MSNs | 251 | 209 | 239 | 98 | 109 |

